# Upregulated extracellular matrix-related genes and impaired synaptic activity in dopaminergic and hippocampal neurons derived from Parkinson’s disease patients with *PINK1* and *PARK2* mutations

**DOI:** 10.1101/2022.12.09.519781

**Authors:** Utkarsh Tripathi, Idan Rosh, Ran Ben Ezer, Ritu Nayak, Ashwani Choudhary, Jose Djamus, Andreea Manole, Henry Haulden, Fred H. Gage, Shani Stern

**Affiliations:** Sagol Department of Neurobiology, University of Haifa, Israel; Laboratory of Genetics, Gage, Salk Institute for Biological Studies, United States; University of Leeds, England

**Author notes:** Equally contributed (Utkarsh Tripathi, Idan Rosh).

## Abstract

Parkinson’s disease (PD) is the second most prevalent neurodegenerative disease. Primary symptoms of PD arise with the loss of dopaminergic (DA) neurons in the Substantia Nigra Pars Compacta, but it affects the hippocampus and cortex also, usually in its later stage. Approximately 15% of PD cases familial with a genetic mutation. Two of the most associated genes with autosomal recessive (AR) early-onset familial PD are *PINK1 and PARK2*. There is a need for in-vitro studies of these genetic mutations in order to understand the neurophysiological changes in patients’ neurons that may contribute to neurodegeneration. In this work, we generated and differentiated DA and hippocampal neurons from iPSCs derived from two patients with a double mutation in their *PINK1 and PARK2* (one homozygous and one heterozygous) genes and assessed their neurophysiology compared to two healthy controls. We showed that the synaptic activity of PD neurons generated from patients with the *PINK1 and PARK2* mutations is impaired in the hippocampus and dopaminergic neurons. Mutant dopaminergic neurons had enhanced excitatory post-synaptic activity. In addition, DA neurons with the homozygous mutation of *PINK1* exhibited more pronounced electrophysiological differences compared to the control neurons. Signaling network analysis of RNA sequencing results revealed that Focal adhesion and ECM receptor pathway were the top 2 upregulated pathways in the mutant PD neurons. These phenotypes are reversed to PD phenotypes of other mutations, suggesting that the interaction of the two mutations may yield different mechanisms of PD.

## Introduction

Parkinson’s disease (PD) is a progressive neurodegenerative disease that is the second most prevalent neurological disease among the elderly^1^. PD affects approximately 2–3% of the world’s population over the age of 65^2^, and it is considered to be one of the aging-related diseases^3–6^. However, genetic mutations can result in an early-onset PD (before the age of 50), also known as familial PD, which accounts for 10-15% of all cases^7^.

Genes involved in monogenic cases of PD include *SNCA*, which codes for the α-synuclein protein, leucine-rich repeat kinase 2 (*LRRK2*), glucosidase beta acid (GBA), phosphatase and tensin homolog (PTEN)-induced putative kinase 1 (*PINK1*), parkin RBR E3 ubiquitin-protein ligase (*PARK2*), and cytoplasmic protein sorting 35 (*VPS35*). The genes *SNCA*, *LRRK2*, and *VPS35* are associated with PD in autosomal dominant forms, whereas *PINK1* and *PARK2* are associated with PD in an autosomal recessive form^8^. Approximately 85% of PD cases are sporadic because no identifiable genetic etiology has been established^9^.

PD is further characterized by the selective loss of the dopaminergic (DA) neurons in the substantia nigra pars compacta (SNpc)^10^. A distinguishing feature of PD is α-synuclein aggregation, which is a primary component of Lewy bodies (LBs) that contributes to the pathophysiology of various PD subtypes^11^. However, there is an ongoing debate about whether the production of α-synuclein aggregations in Lewy bodies is sufficient to cause clinical parkinsonism or neurodegeneration. A recent study has connected PD to an aberrant accumulation of several misfolded proteins, although their specific role in the disease is uncertain. While the precise pathogenic basis of PD remains unclear, several variants with variable prognoses have been extensively characterized^12^.

Although PD is a mobility disorder in some instances, it may also impair cognition and other autonomic processes^13^. The cardinal motor symptoms include tremors, bradykinesia, hypokinesia, akinesia, stiffness, and postural instability. The current state of pharmaceutical treatment does not alter PD’s normal progression. Instead, treatment focuses on symptoms and largely targets the DA system.

*PINK1* and *PARK2* are the most common genes linked to autosomal recessive (AR) early-onset familial PD. Disease onset typically occurs in the third or fourth decade of life and usually presents as a slow progression^14–16^. The *PARK2* gene is located on the 6q25.2–27 chromosome^17^ and encodes the Parkin protein, a cytosolic E3 ubiquitin ligase that functions in the ubiquitin-proteasome pathway. PD onset in people with *PARK2* mutations occurs before the fourth decade of life and constitutes 50% of all AR PD cases. These mutations range from minor deletions and base pair substitutions to massive deletions that span hundreds of nucleotides^18,19^.

Following *PARK2*, the second most frequent AR mutation in young-onset PD is *PINK1*, a serine/threonine protein kinase encoded by the *PINK1* gene. *PINK1* has a kinase domain at the C-terminus and a mitochondrial targeting domain at the N-terminus. The proteasome truncates the N-terminal domain of the gene, resulting in the release of cytosolic *PINK1*^20^. Forty-two different mutations have been documented within the *PINK1* exons in both heterozygous and homozygous states, with Q456X being the most common form of *PINK1*-related PD mutation^21,22^. Recent studies using induced pluripotent stem cells (iPSCs) have shown that *PINK1 and PARK2* mutations are involved in mitochondrial dysfunction and that *PINK1 and PARK2* cooperate to regulate mitophagy, one of the mitochondrial quality control mechanisms^23,24^.

The primary pathogenic symptom of PD is the loss of DA neurons, so the study of PD using iPSCs focuses on DA neuronal differentiation^25,26^. Previous studies that worked on phenotypic characterization using iPSCs became the platform to establish new paradigms for cellular therapeutic techniques, drug development, and preclinical and clinical study screening^27–34^. In iPSC-derived DA neurons, mutant *PARK2* was unable to recruit to *PINK1*, which reduced dopamine vesicle endocytosis, disrupted microtubule stability, decreased neuronal complexity, increased spontaneous excitatory post-synaptic currents in neurons, and increased dopamine release (although there is still no consensus on this specific point). Mutant *PINK1* increased mitochondrial DNA levels, mitochondrial oxidative stress, autophagy, and mitochondrial respiration while interfering with ubiquitin-mediated autophagy in the mitochondria and lysosomes^35,36^.

In this study, we generated and differentiated DA neurons from iPSCs derived from two patients with a double mutation in their *PARK2* (heterozygous) and *PINK1* (one patient with a homozygous mutation and the other with a heterozygous mutation) genes and compared their electrophysiological measures to DA neurons derived from two control iPSC lines. Although PD is considered a movement disorder, many patients also experience non-motor symptoms. Depression and cognitive decline are described in many patients^37^. Changes in the hippocampus have been found that are related to these symptoms^38^. Furthermore, as the hippocampus region is the brain’s stem cell niche, previous data indicate that genetic abnormalities in PD patients may influence adult hippocampal neurogenesis^39^. Therefore, we also differentiated hippocampal neurons from the same patients and the two controls to measure the changes in hippocampal neurons^40^.

We employed electrophysiological techniques to study neurophysiological changes in DA and hippocampal neurons derived from PD patients. We found that *PINK1* and *PARK2* mutant DA neurons showed increased synaptic activity compared to control neurons. This finding was in contrast to all our measurements in DA neurons derived from patients with other mutations and sporadic PD^41^. Interestingly, the patient DA neurons harboring the homozygous *PINK1* mutation (with heterozygous *PARK2*) showed a more severe abnormality than the patient DA neurons harboring the heterozygous *PINK1* mutation (with heterozygous *PARK2*), suggesting the importance of the mutant gene dosage. Additionally, while in our previous study^41^, the reduction in synaptic activity was accompanied by down-regulation of genes related to the brain extracellular matrix (ECM) and focal adhesions, in this study we saw an up-regulation of these pathways, suggesting a connection between the expression of genes from the ECM and focal adhesion pathways and synaptic activity. The hippocampal dentate gyrus (DG) granule neurons derived from the patients with the *PINK1* and *PARK2* mutations showed a different phenotype, suggesting that PD may have a different effect on neurons from different brain areas.

## Material and Methods

### DA neuron differentiation

To generate in vitro midbrain DA neurons, we employed a previously reported protocol^41,42^. Human iPSCs (hiPSCs) obtained from two *PINK1* and *PARK2* patients and two healthy controls were grown until they became ~80% confluent. Accutase and trypsin inhibitor were used to dissociate iPSCs into a single-cell suspension, and the cells were then replated on matrigel-coated plates in mTesR plus media at a density of 40,000 cells/cm2 (Stem Cell Technologies). Cells were allowed to proliferate for two days with a daily media change. When the cells reached 70-80% confluency, the differentiation process was initiated by transferring to KSR media (DMEM F-12 with Glutamax, 15% KO-SR, 1% NEAA, 1% Antibiotic-Antimycotic, 0.1 mM β-mercaptoethanol). The starting date of differentiation was designated as day 0. From day 5 to day 10, the medium was changed gradually to N2 medium (DMEM F-12 with Glutamax, 1% N2 supplement, 1% Antibiotic-Antimycotic) (for day 5 and 6: 75% KSR: 25% N2; whereas for day 7 and 8: 50% KSR: 50% N2; and for day 9 and 10: 25% KSR: 75% N2). The medium was changed to B27 medium on Day 11 (Neurobasal medium, 2% B27 supplement, 1% glutamax, 1% Antibiotic-Antimycotic, 10 ng/mL BDNF, 10 ng/mL GDNF, 1 ng/mL TGFβ3, 0.2 mM ascorbic acid, and 0.1 mM cAMP).

Throughout the differentiation process, small molecule components were introduced to the culture (10 M SB431542 on days 0-4; 100 nM LDN-193189 on days 0-12; 2 M purmorphamine, 0.25 M SAG, 100 ng/mL FGF8b on days 1-6; 3 M CHIR99021 on days 3-12). After 20 to 25 days, neurons were dissociated and replated onto matrigel-coated coverslips and allowed to develop in B27 medium until day 30. On day 30, the base medium was gradually replaced with Brainphys medium to induce synaptic connections. The whole-cell patch-clamp recording was performed on days 45-50.

### Hippocampal differentiation

Additionally, we used a previously published strategy^43,44^ to differentiate iPSCs into hippocampal DG granule neurons. Similar to the differentiation of DA neurons, iPSCs were grown to 80% confluency.

Following this, EBs were created by mechanically dissociating them with dispase, and then they were plated onto low-adherence plates in an mTeSR plus media supplemented with Y-27632 ROCK inhibitor. For 20 days, the EBs were grown in an anti-caudalizing medium comprising DMEM/F12, Glutamax, B27 (without vitamin A), N2, LDN-193189, XAV939, cyclopamine, and SB431542, followed by plating onto polyornithine/laminin (Sigma)-coated dishes in DMEM/F12 (Invitrogen) plus N2, B27 (without vitamin A) and laminin. After a week, rosettes were manually selected based on their morphology, dissociated with Accutase (Chemicon), plated onto poly-l-ornithine/laminin-coated plates, and fed with NPC media made up of the following ingredients: DMEM/F12, Glutamax, B27 (without vitamin A), N2, laminin, and FGF2. To differentiate the cells, ascorbic acid (200 nM), cyclic AMP (cAMP; 500 mg/ml), laminin (1 mg/ml), BDNF (20 ng/ml), and Wnt3a (20 ng/ml) were added to the differentiation medium for 14 days. Three weeks following the onset of the differentiation, the base medium was switched to BrainPhys.

### Immunocytochemistry

Coverslips with DA and hippocampal DG granule neurons were fixed in warmed 4% paraformaldehyde (PFA) for 15 minutes. Following three washes with DPBS, the cells were then blocked and permeabilized in a blocking solution containing DPBS, 0.2% Molecular Grade Triton X-100, and 10% Donor Horse Serum for 60 minutes. Subsequently, we incubated the cells with primary antibodies in the blocking solution at 4°C overnight, using the following antibodies at the stated dilutions for DA neurons: TH (1:500) and MAP2 (1:500), and for hippocampal neurons, PROX1 (1:4000) and MAP2 (1:500). On the next day, the coverslips were washed three times with DPBS (5 minutes each), incubated with the Alexa Fluor™ secondary antibodies and counterstained with DAPI staining solution (1:3000) for 60 minutes at room temperature. Then the coverslips were washed, mounted on slides using Fluoromount-G mounting medium (0100-01, Southern Biotech), and dried overnight in a dark box. The fluorescence signals were detected using a Nikon A1-R confocal microscope, and images were processed using NIS elements 5.21 (Nikon) and microscopy image analysis software Imaris 9.8 (Oxford Instruments).

### RNA extraction, sequencing, and analyses

Total cellular RNA was extracted from 3-5 million DA neurons per sample that were derived from iPSCs of two patients with *PINK1* and *PARK2* mutations and two lines of healthy controls at 7 to 8 weeks post-differentiation using the zymo RNA clean & concentrator kits, according to the manufacturer’s instructions. Further, the RNA was reverse transcribed using the high-capacity cDNA synthesis kit from AB Biosystems. The traditional process for RNA-Sequencing data analysis includes creating FASTQ-format files containing reads sequenced on a next-generation sequencing (NGS) platform. FASTQC 14 v0.11.8 was used to check the quality of the sequenced reads, and STAR (ultrafast universal RNA-seq aligner algorithm) was used to align them to the hg38 human genome. Following this process, each gene’s expression level was evaluated by counting the number of reads aligned to each exon or full-length transcript. The aligned sequences were counted and checked for differential expressions using HTSeq, a Python framework to work with high-throughput sequencing data, and DESeq2, a moderated estimation of fold change and dispersion for RNA-seq data. Protein network path analysis was performed to find significantly differentially expressed genes [false discovery rate (FDR) <0.05] against the KEGG database. The Network Analyst web application was used to create a graphical representation of Protein network path analysis (https://www.networkanalyst.ca). The graph was constructed using nodes and edges. Each node represents either a pathway (large nodes) or genes (small nodes), which are connected by edges (lines). The size of a node increases in proportion to the number of associated genes. Genes are indicated by little red and green circles, with red indicating a significantly up-regulated gene and green indicating a significantly down-regulated gene (FDR<0.05).

### Electrophysiology

Whole-cell patch-clamp recordings were performed on DA neurons derived from two PD patients carrying the *PINK1* and *PARK2* mutations and on DA neurons derived from two healthy controls after 45-50 days of differentiation. Culture coverslips were placed inside a recording chamber that was filled with HEPES-based artificial cerebrospinal fluid (ACSF) containing (in mM) 139 NaCl, 10 HEPES, 4 KCl, 2 CaCl2, 10 D-glucose, and 1MgCl2 (pH 7.5, osmolarity adjusted to 310 mOsm) that had been warmed to 37°C. The recording micropipettes (tip resistance of 10-15 MΩ) were filled with an internal solution containing (in mM) 130 K-gluconate, 6 KCl, 4 NaCl, 10 Na-HEPES, 0.2 K-EGTA, 0.3 GTP, 2 Mg-ATP, 0.2 cAMP, 10 D-glucose, 0.15% biocytin and 0.06% rhodamine (pH 7.5, osmolarity adjusted to 290-300 mOsm). Data were recorded at room temperature using Clampex v11.1 with a sampling rate of 20 kHz.

### Electrophysiology analysis

For hippocampal neurons, we performed whole-cell patch clamp recordings at three different time points: 2-6 weeks (immature neurons, first time point), 6-8 weeks (young neurons, second time point), and 8 −11 weeks (mature neurons, third time point).

### Total evoked action potentials

Neurons were held in current clamp mode at −60 mV with a constant holding current. Following this, current injections were given in 3 pA stages over 400 ms, starting 12 pA below the steady-hold current needed for −60 mV membrane potential. A total of 38 depolarization steps were injected. Neurons that required more than 50 pA to maintain a voltage of −60 mV were excluded from the analysis. The total evoked action potential is the total number of action potentials that were counted in the first 30 (for hippocampal neurons) or 32 (for dopaminergic neurons) depolarization steps in 400 ms recordings.

### Action potential shape analysis

The first evoked action potential (that was generated with the least amount of injected current) was used to examine the action potential shape in terms of spike fast after-hyperpolarization (fAHP), amplitude, width, and threshold. To compute the amplitude of the 5 ms fAHP, the difference between the spike threshold and the membrane potential value 5 ms after the potential returned to pass the threshold value at the end of the action potential, was used. The spike amplitude was determined as the difference between the highest membrane potential during a spike and the threshold. The time it took for the membrane potential to reach 50% of the spike’s amplitude from the ascending to descending portion was used to compute the action potential width (Full Width at Half Maximum). The membrane potential known as the “spike threshold” occurs when a membrane potential depolarizes and its slope sharply increases, producing an action potential (the first maximum in the second derivative of the graph for voltage vs. time).

### Analysis of sodium, fast and slow potassium currents

Current measurements for sodium and potassium were taken in voltage-clamp mode. Voltage steps of 400 ms in the range of −90 to 80 mV were produced while holding the cells at a voltage of −60 mV. The cell capacitance was used to normalize currents. The cell capacitance was acquired through a membrane test of the Clampex SW software. The maximal outgoing current that occurs right after a depolarization step, usually within a time range of a few milliseconds, was used to calculate the fast potassium current. The slow potassium current was taken as the current at the end of the 400 ms depolarization phase.

The sodium and potassium current amplitudes were statistically analyzed at specific test potentials (from −20 to 0 mV for sodium current and from 40 to 80 mV for potassium current). These specific potentials were chosen because they represent the physiological conditions of a healthy neuron.

### Analysis of synaptic activity

In voltage-clamp mode, synaptic activity, i.e., spontaneous excitatory post-synaptic current (EPSC), was recorded. Currents in the patched neurons were monitored while the neurons were maintained at −60 mV. We evaluated the amplitude and rate of synaptic activity using custom-written Matlab code. The cumulative distribution of EPSCs amplitude was calculated for each group. For each cell, the rate of the synaptic activity was calculated by dividing the number of events by the length of the recording (non-active cells were also included, and the event rate was considered 0 for them).

### Calcium imaging experiments and analyses

Calcium imaging was performed at two different time points for hippocampal neurons: initially at 2-3 weeks post-differentiation (first-time point) and later at 6-7 weeks post-differentiation (second-time point).

To observe calcium transients, neuronal cells were incubated for an hour with 2 mM Fluo-5 AM in the ACSF, i.e., recording solution containing (in mM) 139 NaCl, 10 HEPES, 4 KCl, 2 CaCl2, 10 D-glucose, and 1MgCl2 (pH 7.5, osmolarity adjusted to 310 mOsm). After one hour of incubation, the culture coverslips were placed in a recording chamber filled with clean ACSF that had been pre-warmed to 37°C, and imaging of calcium transients was performed with a video microscopy CCD digital camera (SciCam Pro, Scientifica). The images and data were captured and processed using a custom-written GUI in MATLAB 9.8 (R2021a, MathWorks).

A Python script was used to analyze the time series data for the calcium imaging recordings. Calcium transients’ dynamics were examined to differentiate between neurons and astrocytes, which revealed unique activity patterns between the two types of cells. The astrocytes activity was attributed to the slow changes in calcium activity, whereas neurons showed fast transients. To analyze astrocytes and neurons separately, it was necessary to determine if the Region of Interest (ROI) contained an active neuron, an active astrocyte, or an inactive cell. The dynamic differences in fluorescence time series value were used to distinguish astrocytes from neurons.

We typically recorded for 1,800 frames. The frames were collected at ~10 Hz. To check the steepness of the rise of the fluorescence, we calculated the difference between the values of two adjacent frames. The differences Δf(n) between the current and n-6 samples were calculated and normalized by the maximum amplitude obtained over the entire recording period. If the maximum value of normalized Δf (t) exceeded the threshold of 0.05, it was labeled as an active neuron. Signals below the threshold were attributed to inactive cells or astrocytes.

An assessment to determine if ROIs were active astrocytes or inactive cells was performed on those not identified as active neurons. A fluorescence ratio (Δf/f0) was calculated for every signal by dividing the signal’s maximum amplitude by the baseline fluorescence. The lowest value of the signal was attributed as the baseline of the cell. The computed ratio was then compared to the ratio of the most active astrocyte (the astrocyte with the highest fluorescence ratio in a particular recorded area). The signal was considered inactive if its fluorescence ratio was less than 10% of the fluorescence ratio of the most active astrocyte. The remaining signals were characterized as active astrocytes.

### The activity of neurons and astrocytes

A peak detection algorithm was used to count the fast calcium transient events of neurons. The lowest value of the signal was recorded as the baseline of the cell. The average baseline value for all cells was calculated. In addition, the fluorescence of the three most active astrocytes in a recording was averaged. The data were statistically tested and analyzed in Excel using the unpaired Student’s t-test (two-way).

### Correlation calculations

Pairwise correlations were calculated for each ROI (one neuron), with each ROI over 35 samples (approximately 3.5 seconds). A moving average filter was used to calculate these correlations over the sampling time at each sampling point. These correlations were summed for a quantity that represents the correlation of the neuron with its surrounding neurons. The neurons were sorted according to this sum, and the average of the top 10 was obtained as a measure of synchronicity of this area in the recordings.

### The power spectral density (PSD) ratio of all and high frequencies

The PSD is a valuable tool for analyzing the frequencies and amplitudes of oscillatory neurons. After sorting the neurons based on their event counts from selected high synchronized recordings, the PSD was calculated for each neuron and the ratio of area under PSD for the full spectrum and the area under PSD for higher frequency components (greater than 0.1 Hz) was calculated. Then we calculated the mean of this ratio (all frequencies divided by high frequencies) for the top 10 neurons as a measure of this ratio of the area. We compared this ratio of area under the curve for the top most correlated five ROIs and calculated the mean correlation according to these chosen ROIs.

### Statistical analyses

Statistical tests for electrophysiology data were performed using Clampfit-11.1 and Matlab software (R2021a, The MathWorks Inc., Natick, MA, 2000). The p-values were calculated using a two-sample t-test (two-tailed) unless otherwise mentioned (such as the Welch-correction test for comparison of the spike amplitude in DA neurons and Mann-Whitney test for comparing evoked potential in heterozygous and homozygous *PINK1* mutant). The sodium and potassium current changes were analyzed using one-way ANOVA analysis. All data values were presented as mean ± standard error (SE). A value of p<0.05 was considered significant for all statistical tests. For RNA sequencing analysis, the FDR adjusted mean value was evaluated, and a FDR < 0.05 was considered significant.

## Results

### DA neurons from patients with *PINK1* and *PARK2* mutations exhibited alterations in their sodium and potassium currents compared to healthy controls

Our study included two iPSC lines derived from healthy controls and two PD iPSC lines that were derived from patients with *PINK1* and *PARK2* mutations: a 48-year-old patient with a homozygous *PINK1* and heterozygous *PARK2* mutation and a 75-year-old patient with a heterozygous *PINK1* and heterozygous *PARK2* mutation. We differentiated these iPSCs into DA neurons (see Materials and Methods) and examined the intrinsic characteristics and synaptic activity of the neurons using a whole-cell patch clamp.

A total of 45 control and 15 *PINK1* and *PARK2* mutant neurons (eight neurons from the homozygous *PINK1* patient and seven neurons from the heterozygous *PINK1* patient) were patched between 7-10 weeks post-differentiation. Figure 1(o) shows immunostaining with MAP2 (red), which marks mature neurons, and tyrosine hydroxylase (TH green), which marks dopaminergic neurons along with DAPI. Figures 1(a)-(b) show a representative picture of evoked potentials in a control neuron and a *PINK1* and *PARK2* mutant neuron, respectively. The total number of evoked potentials in the first 32 depolarization steps (see Methods) was not significantly different in the *PINK1* and *PARK2* neurons compared to the control neurons (Figure 1(c), p=0.16). However, when we analyzed the recordings from the neurons with the homozygous and heterozygous *PINK1* mutation (and a *PARK2* mutation) separately, we found that the homozygous line had a significantly larger number of evoked potentials compared to the control and heterozygous *PINK1* mutant (Supp. figure 2(a), homozygous *PINK1* mutant vs. control p=0.006, homozygous *PINK1* mutant vs. heterozygous *PINK1* mutant, p=0.043, Using Mann-Whitney test). Figures 1(d) and 1(e) show an example of a single evoked potential of control and *PINK1* and *PARK2* mutant neurons. The fast AHP (5 ms, see Methods) was not significantly different in *PINK1* and *PARK2* mutant neurons in comparison to the control neurons (Figure 1(f), p=0.36). The spike amplitude was significantly lower in *PINK1* and *PARK2* mutant neurons compared to control neurons (Figure 1(g), p=0.038 with Welch-correction test). The spike threshold and cell capacitance were not different between the control neurons and *PINK1* and *PARK2* mutant neurons (Figures 1(h) and 1(i)). Figures 1(j) and 1(k) show examples of the sodium and potassium currents recordings obtained in the voltage-clamp mode of control and *PINK1* and *PARK2* mutant neurons, respectively. The sodium currents (between −20mV to 0mV, see Methods) were significantly decreased (Figure 1(l), p=0.01) in *PINK1* and *PARK2* mutant neurons compared to control neurons. Contrastingly, both the fast potassium and slow potassium currents were found to be significantly larger (between 40 mV to 80 mV) in *PINK1* and *PARK2* mutant neurons compared to the control neurons (Figures 1(m), p=0.01 and 1(n), p=0.047) with 1-way ANOVA analysis.

**Figure 1.**
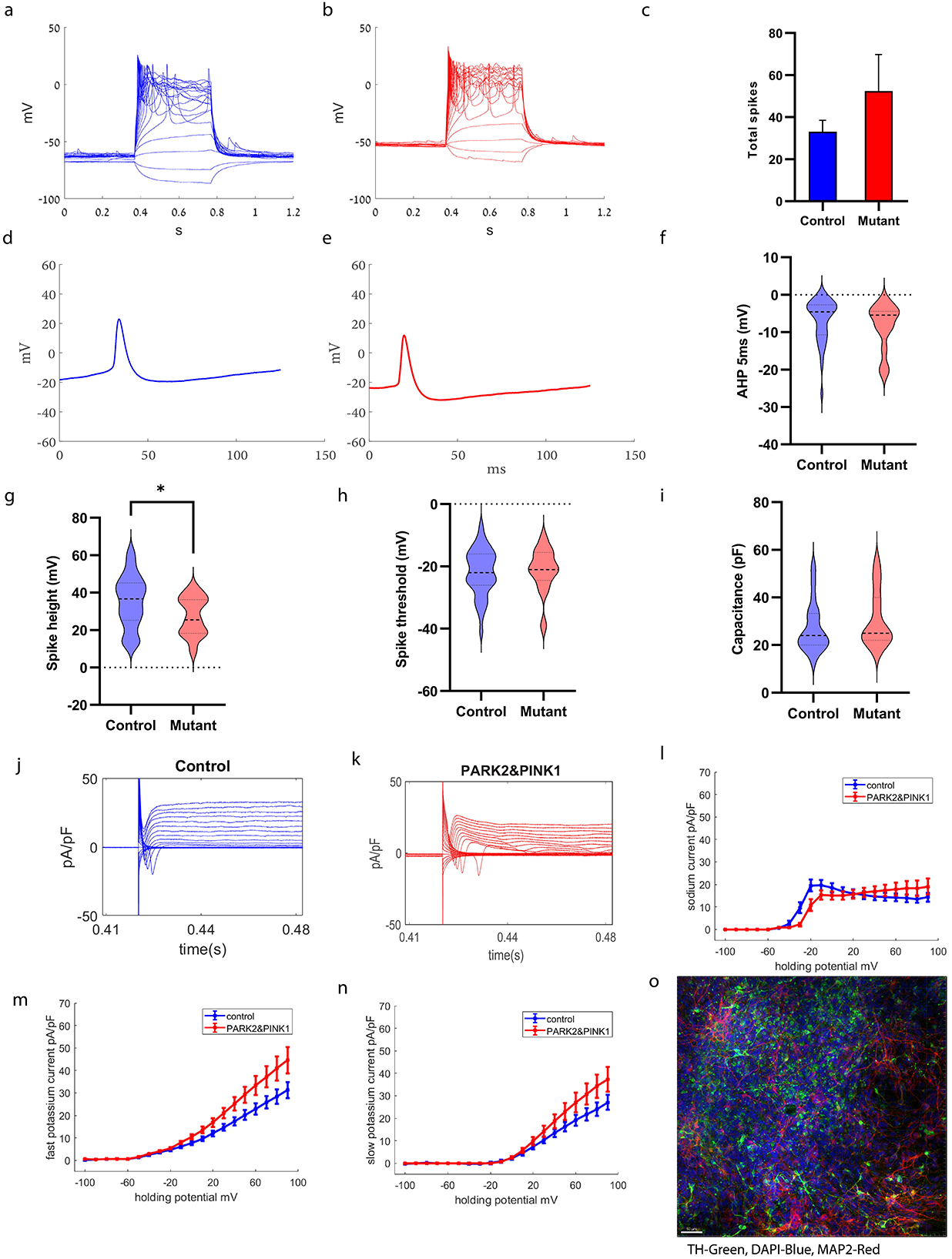
Patient-derived DA neurons exhibited increased potassium currents compared to control neurons. A representative evoked action potential trace in **(a)** control DA neurons and **(b)** DA *PINK1* and *PARK2* mutant neurons. **(c)** The total evoked potential was not significantly different between control and *PINK1* and *PARK2* mutant neurons. A representative picture of the spike shape of **(d)** control and **(e)** *PINK1* and *PARK2* mutant neurons. **(f)** The 5 ms AHP was not different in neurons derived from PD patients with *PINK1* and *PARK2* mutations compared to control neurons. **(g)** The spike height was significantly smaller in neurons derived from PD patients with *PINK1* and *PARK2* mutations compared to healthy controls. Other spike shape features, i.e., **(h)** spike threshold and **(i)** capacitance, were not significantly different between patient-derived DA neurons and control neurons. **j-k**. Representative recordings of sodium and potassium currents in **(j)** control neurons and **(k)** *PINK1* and *PARK2* mutant neurons. **(l)** Sodium current, **(m)** fast potassium currents, and **(n)** slow potassium currents were significantly increased in patient-derived DA neurons using a one-way ANOVA analysis. **(o)** Characterization of DA neurons in culture with immunohistochemistry. TH (green) specifically marks DA neurons in the culture (Scale bar 50 μm). Unless otherwise noted, the error bars in this and the following figures indicate the standard error. Asterisks in this and the subsequent figures denote statistical significance as indicated by the following codes: * p<0.05.

### Increased synaptic activity was observed in DA neurons derived from PD patients with *PINK1* and *PARK2* mutations compared to control neurons

Next, we compared the synaptic activity between control and *PINK1* and *PARK2* mutant neurons. Figures 2(a)-(b) show examples of excitatory post-synaptic currents (EPSCs) in control and *PINK1* and *PARK2* mutant neurons, respectively. The mean amplitude of EPSCs was significantly increased in *PINK1* and *PARK2* mutant neurons compared to the control neurons (Figure 2(c), p=0.0056). In addition, the EPSC rate was significantly increased in *PINK1* and *PARK2* mutant neurons compared to control neurons (Figure 2(d), p=0.001). Further, when we separately analyzed the synaptic activity of the homozygous and heterozygous *PINK1* mutant neurons, we found that the homozygous *PINK1* mutant neurons had an increased EPSC rate compared to control neurons (homozygous *PINK1* mutant vs. control: p<0.0001, homozygous *PINK1* mutant vs. heterozygous *PINK1* mutant: p=0.023, Supp. figure 2(b). The EPSC rate was not significantly different between the heterozygous *PINK1* mutant and control neurons (p=0.74) (Supp. figure 2(b)). Figure 2(e) shows the cumulative distribution of EPSC amplitude of *PINK1* and *PARK2* mutant and control neurons. The cumulative distribution of *PINK1* and *PARK2* mutant neurons’ EPSC amplitude was right-shifted compared to control neurons, indicating higher EPSC amplitudes.

**Figure 2.**
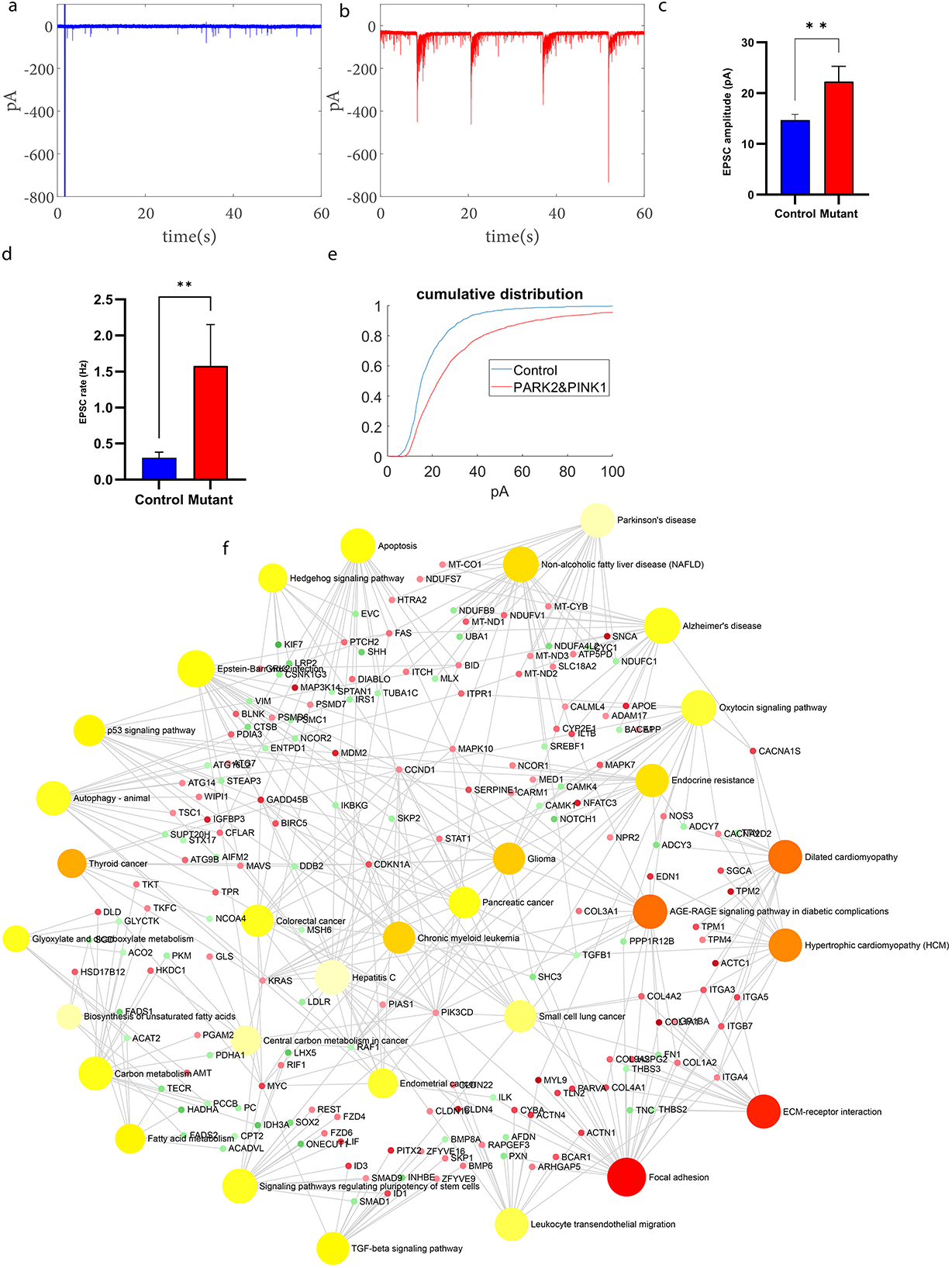
Patient-derived DA neurons exhibited increased synaptic activity compared to control neurons. **a-b**. Example recordings of EPSCs in **(a)** control DA neurons and **(b)** DA *PINK1* and *PARK2* mutant neurons. (**c**) EPSC average amplitude was significantly larger in *PINK1* and *PARK2* mutant neurons compared to control neurons. (**d**) DA *PINK1* and *PARK2* mutant neurons showed significantly increased EPSC rate compared to control neurons. (**e**) The cumulative distribution of EPSC amplitudes was right shifter for *PINK1* and *PARK2* mutant neurons indicating higher EPSC amplitudes compared to control neurons. (**f**) Dysregulated pathways in *PINK1* and *PARK2* mutant DA neurons compared to healthy controls at 7-8 weeks post differentiation. Signaling network analysis with the top enriched KEGG pathways for the *PINK1* and *PARK2* patients’-derived neurons compared to the controls shows that the top dysregulated pathways for this double mutation are “ECM receptor interaction” and “Focal adhesion”. Red circle indicates a significantly up-regulated gene and green indicating a significantly down-regulated gene (FDR<0.05). Asterisks in this and the subsequent figures denote statistical significance as indicated by the following codes: * p<0.05, **p<0.01, *** p<0.001, **** p<0.0001.

### ECM and focal adhesion genes are upregulated in *PINK1* and *PARK2* mutant neurons compared to healthy controls

We next extracted RNA from both the *PINK1* and *PARK2* mutant and control neurons and analyzed gene expression differences. Figure 2(f) presents KEGG signaling pathways that were enriched in neurons derived from patients with *PINK1* and *PARK2* mutation compared to healthy controls. The ECM receptor interaction and focal adhesion pathways were the most dysregulated (upregulated) pathways in *PINK1* and *PARK2* mutant DA neurons compared to healthy controls. The ECM receptor interaction pathway is involved in the regulation of synaptic plasticity^45,46^. The focal adhesion proteins maintain the adhesion of brain cells to the ECM and are essential for neuronal migration and neurogenesis^47,48^.

### Hippocampal neurons derived from patients with the *PINK1* and *PARK2* mutations are hypoexcitable with reduced synaptic activity

We next differentiated hippocampal neurons from the PD patients with the *PINK1* and *PARK2* mutations and performed electrophysiology to compare them to healthy controls. A total of 13 control and 18 *PINK1* and *PARK2* mutant hippocampal neurons were patched at the first time point, 29 control and 20 *PINK1* and *PARK2* mutant hippocampal neurons at the second time point, and 12 control and 10 *PINK1* and *PARK2* mutant hippocampal neurons at the third time point. Figure 3(p) shows immunostaining for hippocampal neurons using MAP2 (red) to mark mature neurons and PROX1 (green) to mark dentate gyrus granule neurons along with DAPI in blue. Figures 3(a)-(b) present a representative image of evoked potentials in control and *PINK1* and *PARK2* mutant hippocampal neurons, respectively, at the second time point. The total number of evoked potentials in the first 30-depolarization steps was not significantly different in control and PINK1 and *PARK2* mutant hippocampal neurons (Figure 3(c)). Figures 3(d)-3(e) show representative traces of EPSCs in control and *PINK1* and *PARK2* mutant hippocampal neurons, respectively. The average amplitude of EPSCs per recorded cell was significantly lower in *PINK1* and *PARK2* mutant hippocampal neurons compared to control neurons (Figure 3(f), p=0.0056). The rate of synaptic events was decreased in *PINK1* and *PARK2* mutants compared to control neurons (Figure 3(g), p=0.0055, using the Mann-Whitney test). The cumulative distribution of the amplitudes of EPSCs in *PINK1* and *PARK2* mutant hippocampal neurons was shifted towards the left compared to control neurons indicating smaller amplitudes (Figure 3(h)). Figures 3(i)-(j) show representative traces of an action potential of control and *PINK1* and *PARK2* mutant hippocampal neurons, respectively. The fAHP (5 ms, see Methods) was not significantly different between control neurons and *PINK1* and *PARK2* mutant hippocampal neurons at the second time point (Figure 3(k)). However, *PINK1* and *PARK2* mutant hippocampal neurons had decreased fAHP compared to control neurons when they were in the early maturation period (first time point) (Supp. figure 1(a)). Figures 3(l)-(m) represent the traces of the sodium and potassium currents acquired in the voltage-clamp mode of control hippocampal and *PINK1* and *PARK2* mutant hippocampal neurons, respectively. *PINK1* and *PARK2* mutant hippocampal neurons that were recorded at the second time point post-differentiation did not show any significant differences in sodium or potassium currents (Figures 3(n) – (o)).

**Figure 3.**
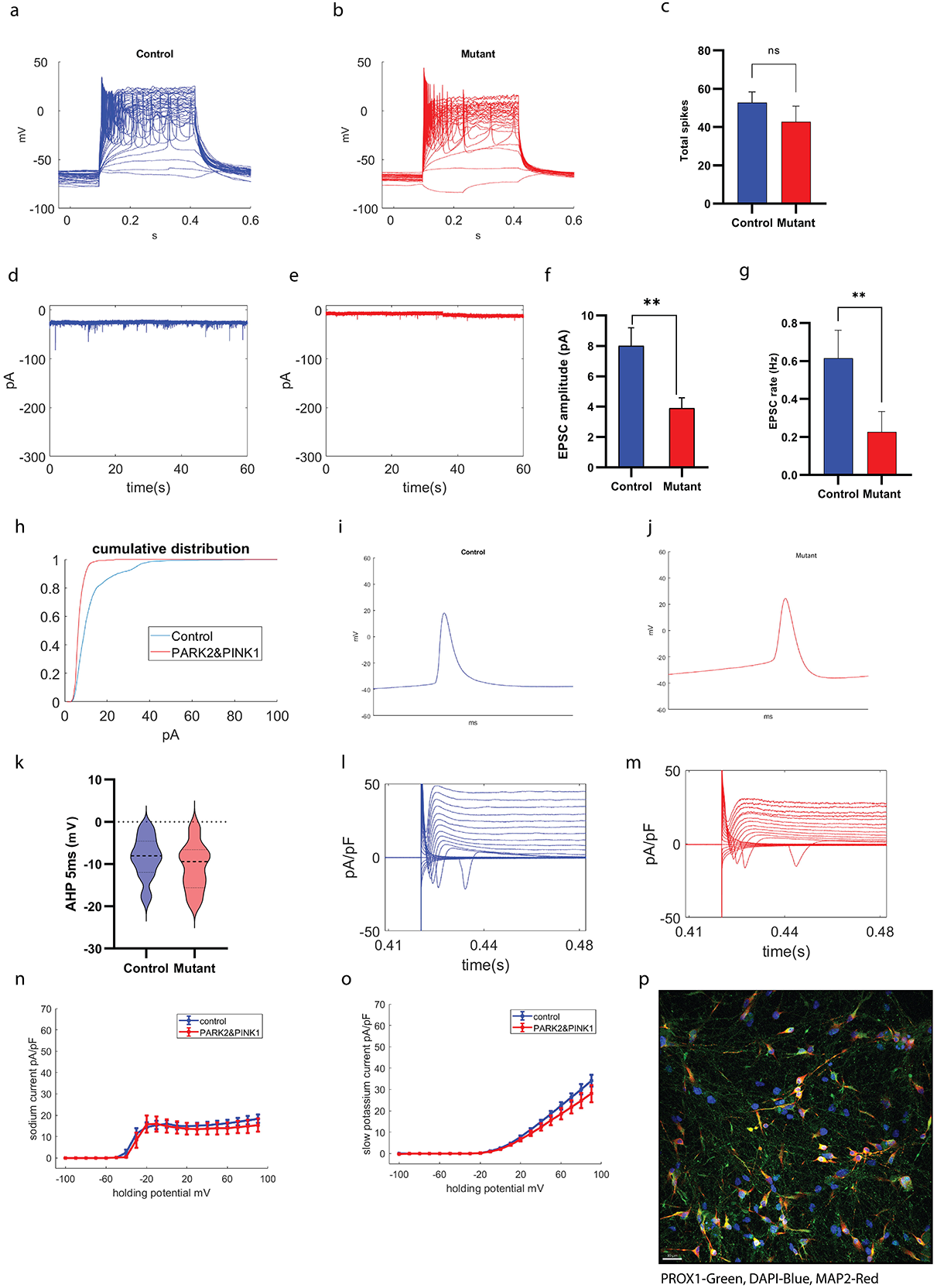
Patient-derived hippocampal neurons possessed reduced excitatory post-synaptic activity compared to healthy controls. Representative evoked action potential traces in (**a**) control hippocampal neurons and (**b**) hippocampal *PINK1* and *PARK2* mutant neurons. (**c**) The total number of evoked potentials was not significantly different between control and *PINK1* and *PARK2* mutant neurons. **d-e**. Example recordings of EPSCs in (**d**) control hippocampal neurons and (**e**) hippocampal *PINK1* and *PARK2* mutant neurons. (**f**) The EPSC amplitude was significantly reduced in *PINK1* and *PARK2* mutant neurons compared to control neurons. (**g**) Hippocampal *PINK1* and *PARK2* mutant neurons showed significantly reduced EPSC rate compared to control neurons. (**h**) The cumulative distribution of EPSC amplitudes showed that EPSC amplitudes of *PINK1* and *PARK2* mutant neurons was left shifted compared to control neurons, indicating smaller EPSC amplitudes in hippocampal *PINK1* and *PARK2* mutant neurons. A representative trace of the spike shape of (**i**) control and (**j**) *PINK1* and *PARK2* mutant neurons. (**k**) The 5 ms AHP was not significantly different in *PINK1* and *PARK2* mutant neurons compared to control neurons. Representative recordings of sodium and potassium currents in (**l**) control neurons and (**m**) *PINK1* and *PARK2* mutant neurons. (**n**) Fast potassium currents and (**o**) slow potassium currents were not significantly different in patient-derived hippocampal neurons compared to controls. (**p**) Characterization of hippocampal neurons in culture with immunohistochemistry. *PROX1* (blue) marks dentate gyrus granule neurons (Scale bar 30 μm). Asterisks in this and the subsequent figures denote statistical significance as indicated by the following codes: **p<0.01.

*PINK1* and *PARK2* mutant hippocampal neurons had slightly reduced slow potassium currents at the first time point (not significant, Supp. figure 1(e)). However, the slow potassium current was significantly increased in *PINK1* and *PARK2* mutant hippocampal neurons compared to control neurons at the third time point (p=0.03, Supp. figure 1(k)). In addition, the sodium current was significantly decreased in *PINK1 and PARK2* mutant hippocampal neurons at the first time point (p=0.0002, Supp. figure 1(f)) but significantly increased in *PINK1* and *PARK2* mutant hippocampal neurons compared to control neurons as they matured (third time point, p=0.0003, Supp. figure 1(l)).

### DG hippocampal neurons derived from patients with the *PARK2* and *PINK1* mutations demonstrated an increased rate of calcium transients compared to control neurons

We also measured the calcium transients (see Methods) in DG hippocampal neurons derived from healthy individuals and *PINK1* and *PARK2* mutant patients. Figure 4(a) shows a fluorescent image of control hippocampal neurons after incubation with fluo-5 calcium indicator. Figure 4(b) shows calcium transient activity (fluorescence intensity in different ROIs) corresponding to the ROIs in Figure 4(a). We observed very synchronous activity between some neurons in a few recordings of heterozygous *PINK1* mutant neurons. To obtain a measure of the connectivity, we calculated the mean correlation (See Methods) between neurons and compared the top five ROIs with the most correlated neurons to compare the heterozygous *PINK1* mutant and control neuronal networks. The mean correlation was significantly increased in heterozygous *PINK1* mutant neurons compared to control neurons (Figure 4(e), p=0.035). Figures 4(c) and 4(d) show an illustration of the correlation heat maps of the most correlated neurons in control and *PINK1* and *PARK2* mutant neuronal networks, respectively. Figures 4(f) and 4(g) represent fluorescent images of control *PINK1* and *PARK2* mutant hippocampal neurons, respectively. These results indicate that, at the second time point, the *PINK1* and *PARK2* mutant neuronal network was more connected and synchronized than the control neuronal network.

**Figure 4.**
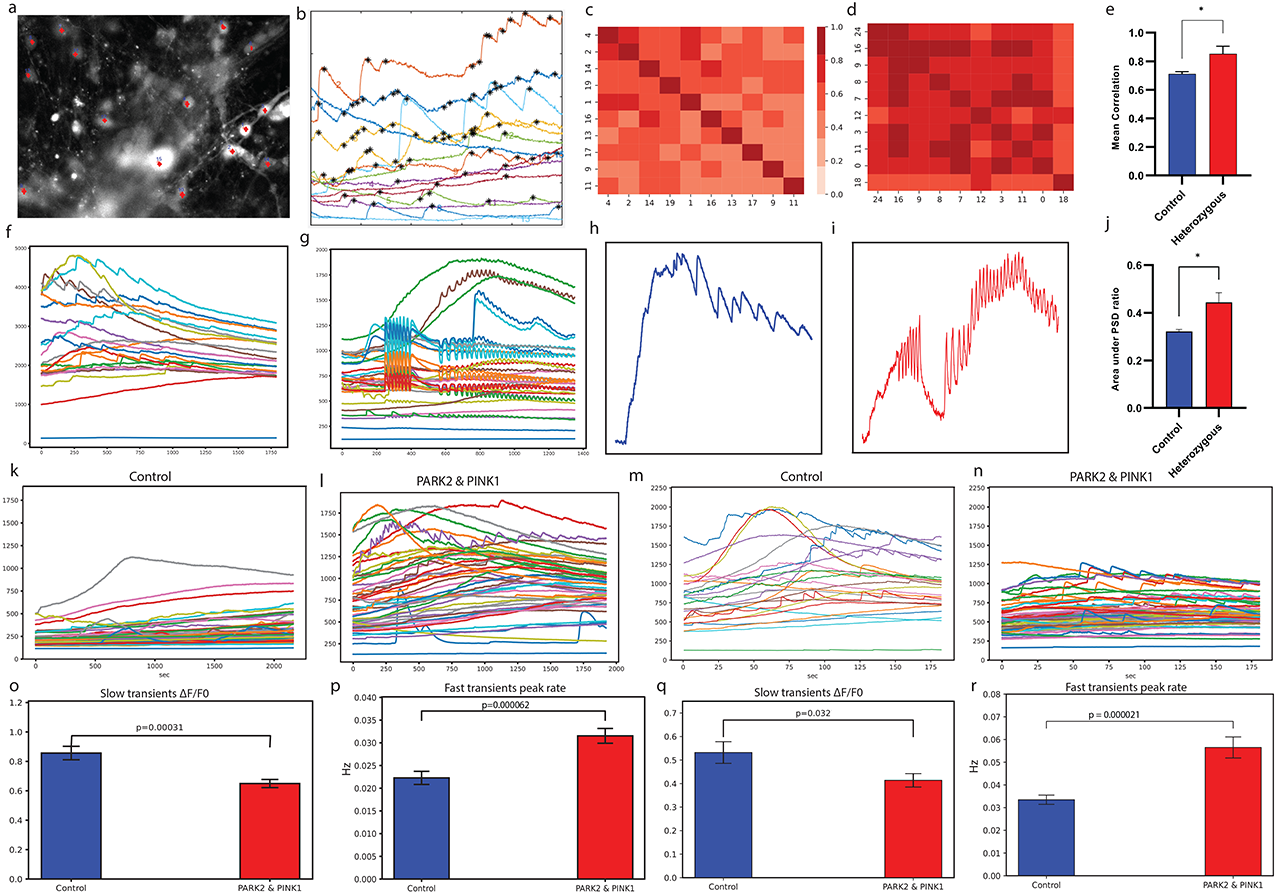
Calcium transient activity was increased and more synchronized in hippocampal neurons harboring *PINK1* and *PARK2* mutations compared to healthy controls. **(a)** A sample video frame of a calcium imaging recording. Red marks represent neurons marked as the region of interest. **(b)** The corresponding fluorescence time series plot for all marked neurons in (a). **c-d**. An example heatmap of the mean correlation value of area recorded area from **(c)** control hippocampal DG granule neuron network and **(d)** hippocampal DG granule neuron network with *PINK1* and *PARK2* mutation. **(e)** The average correlation in the top correlated areas (see Methods) of hippocampal DG granule neurons cultures with *PINK1* and *PARK2* mutations was larger compared to control neuronal culture **f-g**. Representative areas of **(f)** control hippocampal DG granule neurons culture and **(g)** hippocampal DG granule neurons culture with *PINK1* and *PARK2* mutation with high synchronous activity. The *PINK1* and *PARK2* mutant neuronal culture exhibited high synchrony. **h-i**. Representative calcium transient time series plot with a high event frequency of **(h)** a control hippocampal DG granule neuron and **(i)** a hippocampal DG granule neuron with *PINK1* and *PARK2* mutation. The *PINK1* and *PARK2* mutant neuronal culture exhibited high frequency firing patterns. **(j)** The top 5 ratio of area under power spectral density curve of such fluorescent signals between the whole frequency spectrum and higher frequency components was larger in *PINK1* and *PARK2* mutant neurons compared to controls. **k-n**. A representative plot of fluorescence transients at the first time point **(k)** in control neurons and (l) in *PINK1* and *PARK2* mutant neurons. Examples for the second time point are shown in (m) for controls, and (n) for *PINK1* and *PARK2* mutant neurons. Increased activity can be observed in the mutant neurons and higher synchrony can be observed in the mutant neurons in the second time point. **o-p**. At the first time point of calcium imaging **(o)** the average astrocytes fluorescence ratio Δf/fo (see Methods) was significantly decreased in hippocampal *PINK1* and *PARK2* mutant neurons, whereas (**p**) the calcium fast transients’ event (attributed to the neurons) rate was increased in hippocampal *PINK1* and *PARK2* mutant neurons. **q-r**. At the second time point of calcium imaging **(q)** the average astrocytes fluorescence ratio Δf/fo was significantly lower in hippocampal *PINK1* and *PARK2* mutant neurons, and **(r)** the calcium fast transient event rate (attributed to the neurons) was higher in neurons of hippocampal *PINK1* and *PARK2* mutant neurons. Asterisks in this and the subsequent figures denote statistical significance as indicated by the following codes: *p<0.05.

Additionally, we found that some neurons exhibited high-frequency firing in the heterozygous *PINK1* mutant neuronal networks. Therefore, we compared the PSD of the five areas with the highest average frequency in the control and heterozygous *PINK1* mutant neuronal network. Figures 4(h) and 4(i) present control and heterozygous *PINK1* mutant neurons with a high calcium transient event frequency. We calculated the ratio of area under the PSD curves of these fluorescent signals for the entire frequency spectrum divided by the higher frequency component (see Methods). Heterozygous *PINK1* mutant neurons had a significantly higher ratio compared to control neurons (Figure 4(j), p=0.01), which shows that the heterozygous *PINK1* mutant neuronal networks are fast oscillatory behaving neurons compared to the control neuronal network.

We performed calcium imaging at two different time points for hippocampal neurons. Figures 4(k) (m) show illustrative calcium transient time series plots for control and *PINK1* and *PARK2* mutant neurons at two different time points. Figures 4(k) and 4(l) correspond to 2-3 weeks post differentiation. Figures 4(m) and 4(n) correspond to 6-7 weeks post-differentiation. The dynamics of calcium waves in astrocytes are usually reported as being very slow in nature^49–51^. We used these dynamics to classify astrocyte activity in the culture (see Methods). Astrocytes’ average fluorescence ratio (see Methods) was increased for control neurons compared to *PINK1* and *PARK2* mutant neurons during the first time point of calcium imaging (Figure 4(o), p=0.0003) and also at the second time point of calcium imaging (Figure 4(q), p=0.03). The calcium transient event rate in neurons was significantly higher in hippocampal *PINK1* and *PARK2* mutant neurons at the first time point of calcium imaging (Figure 4(p), p=0.00006). *PINK1* and *PARK2* mutant hippocampal neurons exhibited increased calcium transient event rates even after maturation (second time-point of calcium imaging, Figure 4(r), p=0.00002).

## Discussion

Genetic mutations in *PARK2*, *PARK7*, and *PINK1* are among the well-known familial mutations contributing to PD^52^. Mutations in *PINK1* and *PARK2*^36^ have been causally connected to early onset PD. Previous studies have documented the α-synuclein-mediated pathophysiology in PD as a six-stage process^53^. In stage 3 of the pathology, loss of DA neurons in the substantia nigra causes motor symptoms (tremor, bradykinesia, and stiffness) that are evaluated for clinical diagnosis. During stage 4, Lewy pathology affects the hippocampus, causing cognitive dysfunction that progresses to the cortex, causing dementia in later stages^54^. Given this clinical and pathological progression of PD, we investigated both dopaminergic and hippocampal neurons differentiated from PD patient-derived iPSCs. Initial studies on iPSCs of *PINK1* and *PARK2* mutations have found mitochondrial defects and α-synuclein accumulation^24^. Here, we investigated the functional properties of iPSC-derived DA neurons from a patient with homozygous *PINK1* and heterozygous *PARK2* mutation and another patient with heterozygous mutation in the *PINK1* and the *PARK2* genes. The DA neurons of the patients were compared to DA neurons from healthy individuals.

In the combined electrophysiological analysis of DA derived from the PD patients (both homozygous and heterozygous *PINK1* mutant with heterozygous PARK2 mutation), we did not observe a significant difference in the number of evoked action potentials in response to current stimulation compared to healthy controls. Interestingly, when we separated the neurons derived from the patient with the homozygous mutation in the *PINK1* gene, these neurons were significantly hyperexcitable. We further investigated the synaptic activity in *PINK1* and *PARK2* mutant DA neurons and found that the synaptic activity was drastically increased in the *PINK1* and *PARK2* mutant neurons compared to the healthy control neurons. Comparing the synaptic activity between the two *PINK1* and *PARK2* mutant lines showed that the neurons derived from the homozygous patient had a more robust phenotype. Supporting this finding, a previous electrophysiological study of corticostriatal neurons in mice showed differences in synaptic plasticity of corticostriatal neurons with homozygous *PINK1* mutation and a heterozygous mutation^55,56^. In line with these findings, differences in synaptic plasticity (both long-term potentiation as well as long-term depression) were observed during whole-cell recordings in slices of a rodent model of PD with different levels of dopamine denervation, which was induced with different doses of 6-Hydroxydopamine hydrochloride nigrostriatal injection^57^.

Overall, we could detect alterations in electrophysiological properties in *PINK1* and *PARK2* mutant DA neurons that were distinctly stronger in the neurons derived from the patient with the homozygous mutation. A recent study with DA neurons derived from two patients with a *PARK2* mutation showed a reduction in synaptic activity^41^. In the current study, we observed increased synaptic activity, which may indicate some interaction between the two mutations in the *PINK1* and *PARK2* genes and their functional pathways. The high synaptic activity in spiny projection neurons (SPN) has been previously shown in animal PD models^58^. Researchers using mice carrying the knockin mutations *LRRK2-G2019S* or *D2017A* (kinase-dead) showed that, compared to wild-type SPNs, *G2019S* SPNs in postnatal day 21 mice showed a four-fold increase in sEPSC frequency during whole-cell recordings on slices. Through excitotoxicity mechanisms, the high synaptic activity and hyperexcitability of *PINK1* and *PARK2PINK1* mutant DA neurons may contribute to neurodegeneration and synaptic loss. A previous study observed activity-dependent degeneration in GABAergic synapses in the hippocampal culture from presynaptic protein knockout mice^59^. This elevated synaptic activity in dopaminergic *PINK1* and *PARK2* mutant neurons seems paradoxical, as PD is usually associated with synaptic loss^41,60,61^. However, it is possible that biphasic synaptic activity (initially elevated synaptic activity and then decreased synaptic activity at a later time) alterations occur in these mutations just as previously found in Alzheimer’s disease (AD), another disease characterized by severe neurodegeneration^62^, and also in autism models^27,63^.

Furthermore, we performed transcriptomic analysis of the iPSC-derived DA neurons to understand the dysregulated biological pathways and underlying mechanisms related to these double mutations. The ECM receptor interaction and focal adhesion pathways were identified as the most dysregulated in the DA neurons derived from the PD patients with the *PINK1* and *PARK2* mutations. ECM receptor proteins (e.g., integrins) have been reported to play an essential role in developing neuronal structure and signaling mechanisms during synaptic interaction^45^. Both neurodegenerative diseases like AD and neuropsychiatric diseases, including schizophrenia, autism spectrum disorders, and depression, have been linked to ECM protein dysregulation^64–68^. Similarly, focal adhesion proteins are essential in linking ECM proteins to the cytoskeletal elements^69^ and are also known to be critical in AD pathogenesis^70^. These two pathways may interact to regulate crucial signaling mechanisms during synaptic connectivity, which relates to the abnormal synaptic activity observed in *PINK1* and *PARK2* mutant DA neurons. Dysregulation of ECM receptor proteins and focal adhesion proteins has also been reported recently from DA neurons derived from several PD-related mutations and sporadic patients^41^. Interestingly, in that study, ECM and focal adhesion-related genes were down-regulated and the synaptic activity was also decreased, whereas in this study, these pathways are upregulated, yet the synaptic activity is also increased. This finding suggests a link between ECM and focal adhesion pathways and synapse formation.

Although PD is considered a movement disorder, many patients experience non-motor symptoms too. As mentioned above, cognitive decline and depression are among the clinical symptoms observed in PD patients during the later stages of the disease^37^. We found that hippocampal neurons harboring the *PINK1* and *PARK2* mutations have impaired electrophysiological properties, including reduced excitatory synaptic activity compared to the control hippocampal neurons and higher sodium currents in relatively mature neurons compared to the controls. The hippocampus plays a crucial role in cognition, including learning and memory^71^. In mouse models of *PINK1* and *PARK2* mutations, synaptic development and glutamatergic synaptic transmission were impaired in hippocampal neurons^72–74^. These alterations in synaptic activity in hippocampal neurons, as observed in our study, may contribute to the cognitive decline observed in the patients.

Moreover, we also observed that the hippocampal neurons derived from the patients with the *PINK1* and *PARK2* mutations had more pronounced calcium transients. Aging has been shown to increase the calcium transients in hippocampal neurons in rodent models^75,76^. Other studies also suggested changes in calcium homeostasis in hippocampal neurons that progress with aging, leading to alterations in synaptic strength (long-term depression and long-term potentiation)^77^. Evidence of elevated calcium signaling has also been demonstrated in mouse models of AD, resulting in neurite degeneration^78^. Furthermore, several studies have reported the biological role of *PINK1* and *PARK2* genes in mitochondrial biogenesis and trafficking^66,67^ and the alteration in mitochondrial structure in iPSC-derived neurons with *PINK1* and *PARK2* mutations^24^. Thus, we speculate that this alteration in calcium transients observed in a hippocampal neuron could be partly due to the effect of the mutation of *PINK1* and *PARK2* on mitochondria, as mitochondria play a crucial role in regulating calcium homeostasis^78^. Increased calcium has been suggested to activate the calpain-mediated signaling pathways, which could affect learning and memory^77^. This is also likely to indicate functional changes in hippocampal neurons before the appearance of neurodegeneration.

In conclusion, we conducted the first detailed functional study of dopaminergic and hippocampal neurons in iPSC-derived neurons from PD patients with *PINK1* and *PARK2* double mutations and found alterations in electrophysiological properties as well as calcium homeostasis. These findings are also consistent with clinical symptoms found in PD patients and provide an exciting opportunity to use such iPSC-based PD models as a preclinical drug-screening platform.

## Acknowledgments

The authors would like to thank the Israel Science Foundation (ISF grant 1994/21 and 3252/21) and Zuckerman (Zuckerman STEM leadership program) to Dr. Shani Stern and the JPB foundation to Prof. Fred Gage for funding and support.

---

**Supplementary figure 1.**
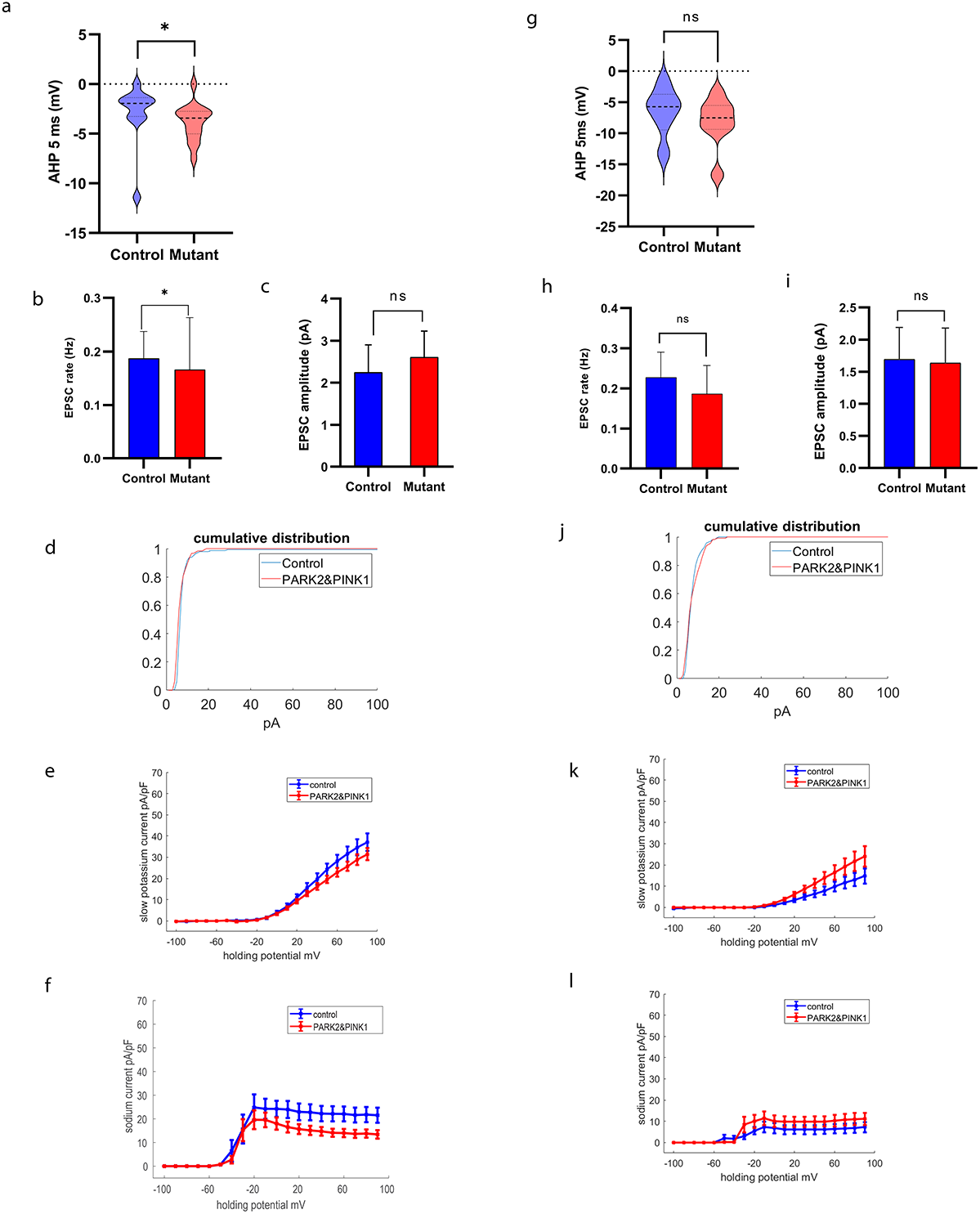
Hippocampal *PINK1* and *PARK2* mutant neuron’s synaptic activity was not different compared to controls during early and post-maturation periods. **a-f**. During the 2-4 weeks post differentiation period (first time period) of hippocampal neurons **(a)** the 5 ms AHP was significantly reduced in *PINK1* and *PARK2* mutant neurons. **(b)** The EPSC rate was decreased in *PINK1* and *PARK2* mutant neurons compared to healthy controls. **(c)** The EPSC amplitude was not significantly different between *PINK1* and *PARK2* mutant neurons and control neurons. **(d)** The cumulative distribution of EPSC amplitude showed that EPSC amplitude distribution was similar for *PINK1* and *PARK2* mutant and control neurons. Initially, **(e)** the sodium and **(f)** the slow potassium currents were significantly decreased in *PINK1* and *PARK2* mutant neurons compared to control neurons. **g-l**. During the 8-11 weeks post differentiation period (the third time point) in hippocampal neurons **(g) the** 5 ms AHP was not significantly different in *PINK1* and *PARK2* mutant neurons**. (h)** The EPSC rate was not significantly different between *PINK1* and *PARK2* mutant and control neurons. **(i)** Also the EPSC amplitude was not significantly different between *PINK1* and *PARK2* mutant and control neurons. **(j)** The cumulative distribution of EPSC amplitudes showed that EPSC amplitudes were similar for *PINK1* and *PARK2* mutant and control neurons. In the later phase (third time point), **(k)** the sodium and **(l)** the slow potassium currents became significantly larger in *PINK1* and *PARK2* mutant neurons compared to control neurons. Asterisks in this and the subsequent figures denote statistical significance as indicated by the following codes: * p<0.05.

**Supplementary figure 2.**
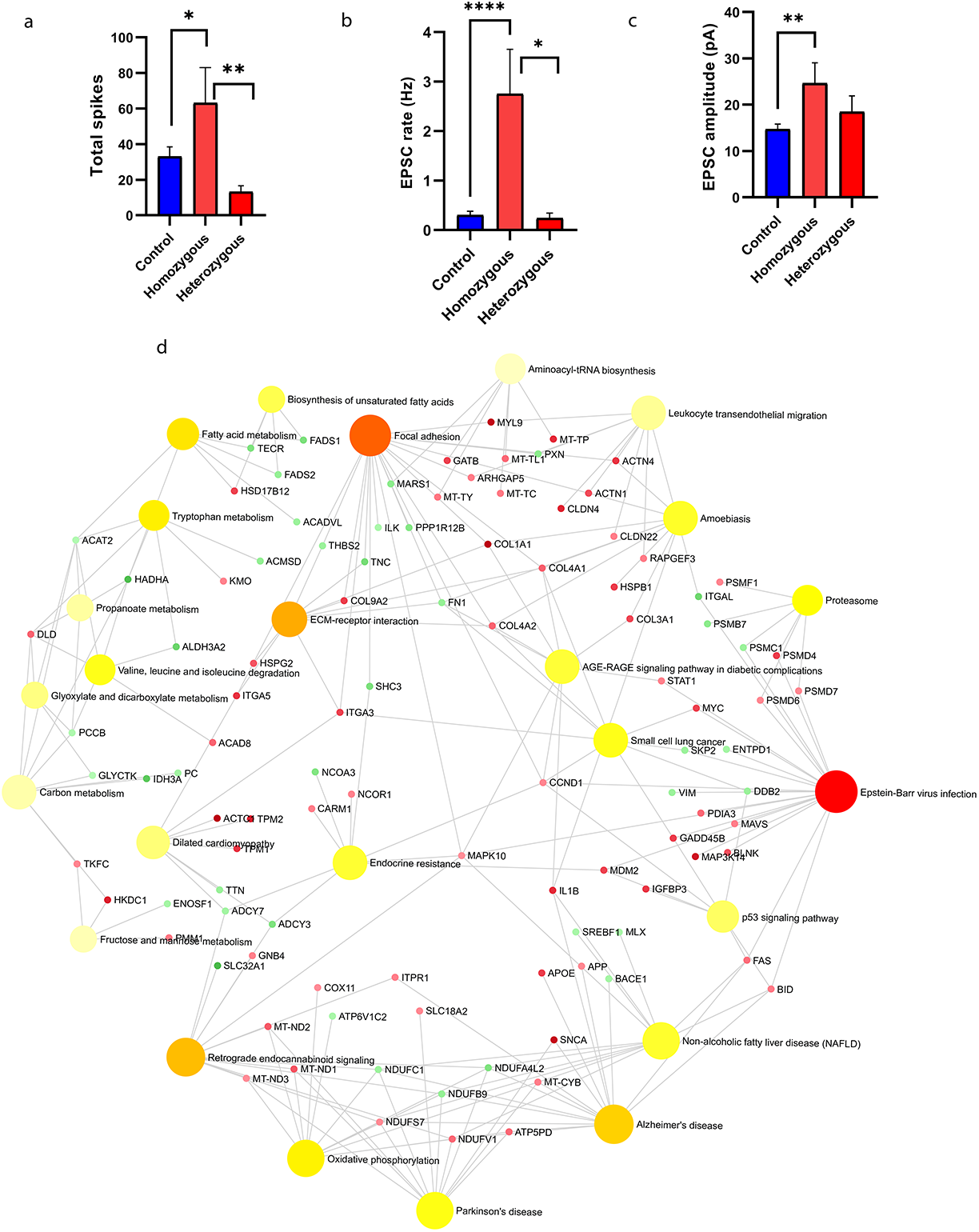
DA neurons derived from patients with homozygous *PINK1* and *PARK2* mutations showed a strong increase in synaptic activity. **(a)** The total evoked number of spikes in DA neurons with homozygous *PINK1* and *PARK2* mutations were significantly increased compared to control neurons and neurons with heterozygous *PINK1* and *PARK2* mutations. **(b) The** EPSC rate in DA neurons with the homozygous *PINK1* mutation was significantly increased compared to control neurons and neurons from the heterozygous *PINK1* mutant patient with the heterozygous *PINK1* mutation. **(c)** Also, the EPSC amplitude was significantly increased in the homozygous *PINK1* mutant neurons compared to the control and the heterozygous *PINK1* mutant line. **(d)** Signaling network analysis with the top enriched KEGG pathways for the homozygous *PINK1* and *PARK2* patient’s-derived neurons compared to the controls shows that the top dysregulated pathways for this double mutation are “Epstein Barr virus infection", “ECM receptor interaction” and “Focal adhesion”.

